# Multilocus microsatellite typing (MLMT) reveals high genetic diversity of *Leishmania infantum* strains causing tegumentary leishmaniasis in northern Italy

**DOI:** 10.64898/2026.02.24.707666

**Authors:** Gianluca Rugna, Elena Carra, Mattia Calzolari, Federica Bergamini, Alice Rabitti, Tommaso Gritti, Margherita Ortalli, Tiziana Lazzarotto, Valeria Gaspari, Germano Castelli, Federica Bruno, Gerald Frank Späth, the Skin_Leish_RER_network, Stefania Varani

## Abstract

**Background:** Tegumentary leishmaniasis (TL) caused by *Leishmania infantum* has re-emerged in northern Italy, raising questions about the genetic diversity and population structure of circulating parasites and their potential role in shaping different clinical outcomes.

**Methodology/Principal findings:** Multilocus microsatellite typing (MLMT) based on 15 polymorphic loci was applied to 44 *L. infantum* strains obtained from TL cases diagnosed between 2013 and 2024 in the Emilia-Romagna region. These strains were compared with sympatric isolates from VL cases, dogs and sand flies. MLMT revealed a considerable genetic variation among TL-associated strains, with 43 distinct multilocus genotypes identified. Population structure analyses using Bayesian clustering, multivariate approaches and phylogenetic reconstruction consistently identified three highly differentiated genetic populations (F_st_ >0.25). TL strains were divided into two main populations: one shared with VL-associated strains (PopB; 9/44) and a second population found exclusively among TL cases (PopC; 28/44). In contrast, the canine-associated population (PopA) showed no overlap with TL cases in this region. Populations also displayed divergent heterozygosity patterns, as indicated by positive and negative Fis values.

**Conclusions/Significance:** These findings revealed previously unknown diversity within *L. infantum* in the study area and demonstrated that inclusion of tegumentary strains is essential to uncover hidden components of parasite population structure. The identification of a TL-associated population supports the existence of multiple evolutionary pathways and emphasises the importance of integrated One Health surveillance, which combines data from humans, animal hosts and vectors to improve understanding of the epidemiology of leishmaniasis in Italy.

**Author summary:** *Leishmania infantum* is a parasite transmitted to humans through the bite of infected insect vectors. It can cause different forms of leishmaniasis, ranging from a systemic disease known as visceral leishmaniasis to a less common form that affects the skin and mucous membranes, called tegumentary leishmaniasis. Dogs are the main reservoir of the parasite and play a key role in maintaining its circulation in endemic areas. In recent years, cases of tegumentary leishmaniasis have re-emerged in northern Italy. This unexpected increase has raised questions about how the parasite is spreading and whether genetic differences among the parasites could explain these new patterns. To explore this, we examined parasitic DNA obtained from tegumentary leishmaniasis cases and compared it with DNA from patients with visceral leishmaniasis, from dogs and insect vectors from the same area. By examining multiple genetic markers, we found that parasites causing the tegumentary form of the disease are genetically diverse and belong to different groups. Notably, one parasite group was found only in cases of tegumentary leishmaniasis and not in visceral infections nor in infected dogs, suggesting that some parasite lineages may be more closely associated with skin and mucosal disease. Overall, our findings show that studying parasites from cutaneous and mucosal lesions provides important information that would otherwise remain hidden. By combining data from humans, animals and insect vectors, our study highlights the importance of integrated surveillance systems for improving our understanding of disease spread and supporting effective public health strategies.

## Introduction

Leishmaniasis is a vector-borne disease caused by hemoflagellate protozoa of the genus *Leishmania* that are transmitted through the bite of infected sand flies belonging to the Phlebotominae subfamily (1). Around 20 of more than 50 *Leishmania* species can cause a wide spectrum of human disease.

The *Leishmania* parasite exhibits a dixenous life cycle, alternating between the promastigote stage in the sand fly vector and the amastigote stage within macrophages of mammals (2). Although all *Leishmania* species share a similar life cycle and are transmitted through the skin, they exhibit varying affinities for specific host tissues (3). Dermotropic species such as *L. tropica* and *L. major* in the Old World often remain confined to the skin, leading to localized infections that manifest as chronic skin lesions, characteristic of cutaneous leishmaniasis (CL). In contrast, viscerotropic parasites of the *L. donovani* complex, such as *L. donovani* and *L. infantum,* disseminate throughout the body and cause a life-threatening visceral leishmaniasis (VL). In VL, amastigote-infected monocytes and macrophages disseminate the parasite through the bloodstream to the spleen, liver and bone marrow, causing the activation of the mononuclear phagocyte system and leading to prolonged fever, splenomegaly, hepatomegaly and pancytopenia (4). Despite being predominantly considered viscerotropic species, increasing evidence indicates that species within the *L. donovani* complex possess dermotropic variants with a wide geographic distribution (5).

In southern Europe, leishmaniasis is mainly caused by *L. infantum* (6). In humans, *L. infantum* has been mostly associated with VL (7), but can also cause localized chronic disease, with parasites replicating in skin macrophages (CL) or in the mucous membrane of the upper respiratory tract (mucosal leishmaniasis, ML). CL and ML are grouped under the term tegumentary leishmaniasis (TL). Within the southern European epidemiological context, the dual tropism of *L. infantum* is attracting increasing attention.

Genetic differences within *L. infantum* strains could influence disease pathology; variation in species-specific genes, gene polymorphisms, pseudogenes, and expression levels of virulence and stress response genes are believed to contribute to the wide range of clinical manifestations (7). However, knowledge on diversity between dermotropic and viscerotropic strains within the *L. infantum* species remains limited. Additional crucial factors influencing *Leishmania* tissue tropism include parasite-vector interactions, vector components such as saliva, and mammalian host factors, including host immunity (2,5). Therefore, the link between parasite diversity and disease outcome is complex and incompletely understood.

In the last fifteen years, human leishmaniasis has emerged in Bologna and Modena provinces, northern Italy, as well as in other areas of the same region (Emilia-Romagna region, E-R) (8–10). Interestingly, the *Leishmania* strains isolated from this epidemic focus possess genetic features that diverge from the main *L. infantum* population defined as Linf1/MON-1 (11–15). Next generation sequencing (NGS) analysis revealed that VL strains from this geographic area exhibit features typical of *L. infantum/L. donovani* hybrids (13). In addition, by typing two *Leishmania* genomic targets that are useful for species identification, i.e. the heat shock protein 70 *(Hsp70*) and the cysteine peptidase b *(Cpb*), we found that TL strains from the same area possess genetic signatures of both *L. infantum* and *L. donovani* (12,14), thus suggesting a hybrid nature also for the dermotropic strains.

Among the genotyping methods investigating the population genetics of the *L. donovani* complex, multilocus microsatellite typing (MLMT), which relies on length variations of microsatellite markers, has been shown to effectively discriminate parasitic strains and has emerged as a valuable molecular tool for elucidating the epidemiology of leishmaniasis and the population dynamics of the parasite (16) (17). This discriminatory power arises from the highly polymorphic and co-dominant nature of microsatellite markers, which are distributed in different chromosomes of the parasite.

In this study, a panel of microsatellite markers was employed to investigate the genetic structure and diversity of *L. infantum* strains obtained from TL cases in northern Italy. The resulting genetic profiles were compared with those of previously published *L. infantum* strains from humans (VL cases), dogs and sand flies from the same geographic area to assess their genetic relationship.

## Methods

### Origin of isolates and biological samples

MLMT analysis was performed *de novo* on TL samples, as well as on selected canine and sand fly strains that were analyzed specifically for this study. These newly generated data were then compared with previously published MLMT datasets of *L. infantum* from our previous studies, which included human cases of VL, isolates obtained from canine leishmaniasis (CanL) cases, as well as sand fly strains (13,15). The TL specimens that were typed by MLMT in this study were in total *n* = 44; of these *n* = 24 thin sections from formalin-fixed and paraffin embedded biopsies (FFPE), *n* = 16 fresh specimens from punch biopsies, and four *Leishmania* isolates (MHOM/IT/2021/IZSLER-BO001, MHOM/IT/2023/IZSLER-BO003, MHOM/IT2023/IZSLER-BO004, MHOM/IT2024/IZSLER-BO005) obtained from skin/mucosal lesions of TL cases. Parasite species of TL samples was defined through amplification and sequencing of the ribosomal internal transcribed spacer 1 (ITS)-1 fragment of 320 bp or of the *Hsp70* coding sequence fragment of 262 bp as described (12).

Canine strains were obtained from dogs with CanL at the Experimental Zooprophylactic Institute of Lombardy and Emilia Romagna (IZSLER), Modena, Italy. Sand fly samples were collected by IZSLER in the frame of the Regional surveillance plan for canine leishmaniasis (18). Samples were stored at -80 °C at the Unit of Microbiology (University Hospital of Bologna, Italy) for human strains, while storing of CanL isolates and sand fly strains, and MLMT evaluation were performed at the IZSLER, Modena Unit (Italy).

In total, we analyzed 133 *L. infantum* strains. These included (i) 44 TL strains described in the present study, (ii) 52 canine isolates, (iii) 12 sand fly strains, and (iv) 25 human (VL) strains from E-R. MLMT data were already available for 73 strains (25 human VL, 40 canine, and 8 sand fly strains) from a previous study (13), while 16 additional isolates (12 from dogs and 4 from sand flies) were newly analyzed for this work to reduce potential bias due to sampling year or location. The complete set of MLMT data generated and/or analyzed in this study is available in S1 Table.

### DNA extraction

DNA extraction was performed using standardized commercial kits. For fresh tissue samples obtained from punch biopsies, DNA was extracted with the DNeasy Blood and Tissue Kit (Qiagen, Hilden, Germany) according to the manufacturer’s instructions. The same kit was also used for human, canine, and sand fly isolates. For FFPE specimens from TL cases, DNA extraction was carried out using the Maxwell® CSC DNA FFPE Kit (Promega, Madison, WI, USA) and the Maxwell CSC instrument, following the manufacturer’s protocol.

### Multilocus microsatellite typing (MLMT)

A panel of 15 dinucleotide microsatellite markers, previously validated for the *L. donovani* species complex, were used to determine genetic diversity indexes in the *L. infantum* strains. The markers included Li41-56, Li46-67, Li21-34, Li22-35, Li23-41, Lm2TG, Lm4TA, Li71-5/2, LIST7039, Li71-33, Li71-7, CS20, Li45-24, TubCA and LIST7031. The amplification reaction of the microsatellite loci was performed as previously described (15) using fluorescence-labeled forward primers (WellRed dyes, Sigma-Aldrich, Saint Louis, USA). The amplicons were subjected to automated fragment analysis on the capillary Sciex GenomeLab™ GeXP sequencer (ex-Beckman Coulter) (MA, USA) and the fragment sizes were determined using the “Fragment” package of the GenomeLab™ System software, version 11.0.24.

The microsatellite repeat numbers were calculated for all loci by comparing the sizes of the respective fragments to those of the strain MHOM/TN/1980/IPT1, which was included in each run as a standard for fragment size, and for which the repeat numbers had been determined by direct sequencing (as described in supplementary protocol in (15)). The repeat numbers for all markers were then assembled into a multilocus genotype (MLG) for each sample under study.

### Population structure

The 44 TL strains typed in this study were first analysed to assess microsatellite diversity and MLG distribution. For population structure inference, these strains were combined with previously characterized strains, resulting in an integrated dataset of 133 strains. To minimize bias due to clonal redundancy, clone-correction was applied by retaining each unique MLG only once. The clone-corrected dataset (n=106) was used for Bayesian clustering and population genetic analyses. The MLG data were analyzed using both a Bayesian model-based approach implemented in STRUCTURE v.2.3.4. software package (19) and a discriminant analysis of principal components (DAPC), which is a model-free clustering method (20).

STRUCTURE determines genetically distinct populations based on allele frequencies and estimates the individual’s membership coefficient (Q-value) in each probabilistic population. The analysis was run using 500,000 Markov chain Monte Carlo iterations after 50,000 burn-in generations under the admixture model and correlated allele frequencies. Ten independent simulations were performed for each value of K (estimated number of populations) between K=1 and K=10. The best-fitting value of K was chosen by calculating ΔK statistics (21). To resolve label switching, results were aligned with CLUMPP v1.1.2 (22) and mean membership coefficients (Q) were computed across runs. Strains were assigned to populations according to their posterior membership coefficients (Q-values), following a scheme adapted from Cortes et al. (23) for K=3. Strains with Q1 > 0.75 were classified as belonging to ‘core’ population (e.g., population A, PopA). If 0.60 ≤ Q1 ≤ 0.75 and Q2 ≥ 0.10, they were considered admixed between two populations with a predominance of one of them (e.g., PopAA/B or PopA/BB). Strains with 0.40 ≤ Q1 < 0.60 were assigned as admixed between two (e.g., PopA/B or PopB/C) or three populations if Q3 ≥ 0.10 (e.g., PopA/B/C). Following STRUCTURE analysis on the clone-corrected dataset, clonal replicates were reintroduced and assigned to the same population as their corresponding MLG for descriptive purposes.

STRUCTURE results were compared with a discriminant analysis of principal components (DAPC), implemented in the *adegenet* v2.1.11 package (24,25) in RStudio v2025.05.1 (26). DAPC is a multivariate method that does not rely on predefined population genetic models. It first transforms multilocus genotypes using principal component analysis (PCA) to reduce dimensionality and subsequently performs discriminant analysis to maximize between-group variation while minimizing within-group variation. DAPC was performed on the clone-corrected dataset, and concordance with STRUCTURE-inferred populations was assessed. The most likely number of clusters was inferred using the Bayesian information criterion (BIC) implemented in the function *find.clusters*. The optimal number of retained principal components was determined by cross validation (CV). All plots were generated in RStudio using the ggplot2 package (27).

### Population genetics characterization of inferred populations

Descriptive statistics for microsatellite markers and genetic populations were calculated using the GDA software v1 (28). These analyses included allelic diversity (number of allelic variants per marker, A, and mean number of alleles per population, MNA), proportion of polymorphic loci (P), and expected (He) and observed (Ho) heterozygosity. Genepop on the Web v4.7.5 (29) was used to test for Hardy-Weinberg equilibrium (HWE) and to estimate the inbreeding coefficient (F_is_) across all locus/population combinations by means of Markov Chain permutations (30) with 1000 dememorization steps, 400 batches, and 1000 iterations per batch. Fisher’s combined probability test was used to calculate *p*-values across loci and populations, with and without correction for multiple testing using the Bonferroni method. Genetic differentiation between populations was further assessed using F-statistics by estimating F_st_ (fixation index), which compares genetic variation within populations to genetic variation between populations. F_st_ values were calculated using the Weir and Cockerham estimator (31) as implemented in the *hierfstat* package v0.5-11 (21) in RStudio. F_st_ values higher than 0.25 with significant *p*-values (<0.001) were considered to indicate strong genetic differentiation (32).

### Microsatellite-based tree analysis

To visualize genetic relationships among strains and assess concordance with STRUCTURE and DAPC results, a microsatellite-based tree was inferred using the BEASTvntr package implemented in the BEAST2. 2.4.3 (33). The diploid genotypes were entered as two distinct partitions, with a linked tree and strict clock. The Sainudiin microsatellite mutation model (34) was applied, and analyses were run for ten million MCMC steps. The tree was inferred from the integrated dataset of 133 strains and included selected WHO reference strains (S1 Table), representative of *L. infantum* zymodemes MON-1, MON-72, MON-24, as well as the *L. donovani* strain MHOM/NP/02/BPK282/0cl4, which were included exclusively as external references to anchor major clades; three profiles were retrieved from previously published datasets (13), while one profile was generated in the present study and used only for evolutionary comparison. Trees were visualized with iTOL v6 (https://itol.embl.de/).

### Ethical clearance

*Leishmania* samples were obtained from TL cases within the diagnostic and surveillance activities of the Regional Reference Laboratory for human leishmaniasis (RRL) at the Unit of Microbiology, University Hospital of Bologna, Italy in the period 2013-2024. Samples were coded and anonymized. The study was conducted in accordance with the declaration of Helsinki, and the protocol was approved by the Ethics Committee of the Area Vasta Emilia Centro (study number: n° EM414-2023_97/2017/O/Tess/AOUBo). Written informed consent was obtained from patients included in this study.

All animal samples were obtained for diagnostic purpose with no unnecessary invasive procedures, as part of the E-R surveillance program on leishmaniasis, including parasitological confirmation of CanL in clinically suspected dogs. Evaluation of these samples did not require ethical approval according to the European Directive 2010/63/EU. Oral informed consent was obtained from the owners of dogs at the time of clinical examination.

## Results

### Samples’ characteristics

A total of 44 samples were typed for this study; 39 cases were CL, while the remaining five cases were ML. Thirty-seven samples were obtained from TL cases residing in five provinces of E-R, of these 23 (62%) were from the Bologna province (Table 1). The remaining samples were obtained from TL cases likely infected in different Italian regions (three cases) or for which the place of infection was unknown (four cases). The designation and species typing results of all TL samples included in the study are listed in S1 Table. The sequencing of the ITS1 or *Hsp70* PCR products identified all the samples as *L*. *infantum* (see accession number, A.N., in S1 Table).

**Table 1.**
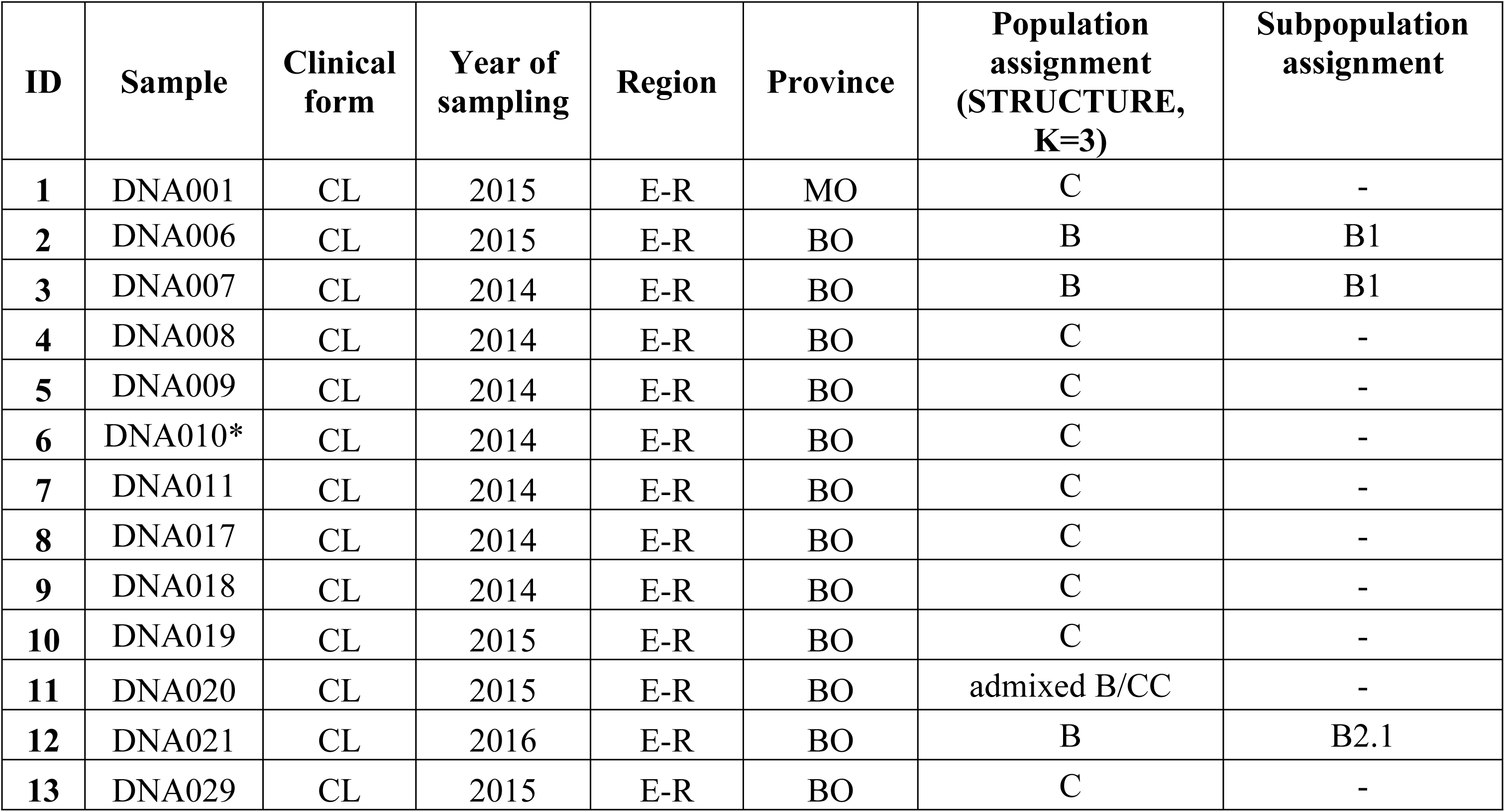
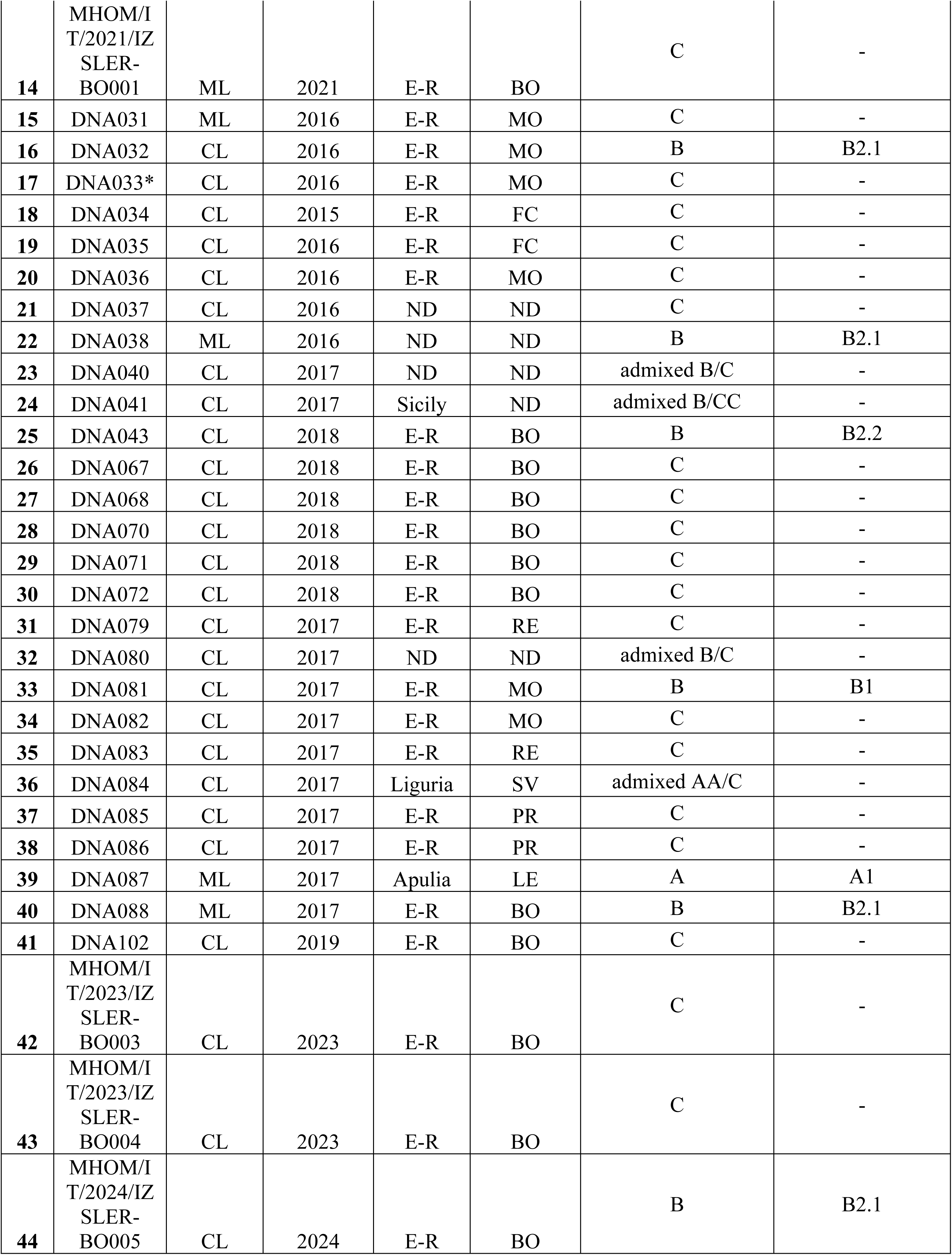
Characteristics of tegumentary leishmaniasis (TL) samples included in the study and population assignment.

### Microsatellite variation and genetic diversity in dermotropic *L. infantum* strains

Fifteen polymorphic microsatellite loci were used to assess the genetic variation of the 44 *L. infantum* samples obtained from TL cases diagnosed in E-R. Amplification reactions were successfully performed for 42 samples, while the sample ID34 resulted negative for the loci Li22-35 and Li45-24, and the sample ID23 resulted negative for the loci Li71-33, Li45-24 and List7031. As shown in Table 2, all markers were polymorphic, exhibiting between 3 and 11 different alleles per marker. The most variable markers were Lm4TA and Lm2TG (11 and 10 alleles, respectively), while Li46-67 was the least polymorphic, presenting only 3 alleles. The mean number of allelic variants per locus (A) was 6.3. The observed heterozygosity (Ho) was seen in 14 out of 15 loci and was between 0.205 and 0.614. The expected heterozygosity (He), representing the probability that an individual will be heterozygous over the loci tested, ranged from 0.403 to 0.821 and it was in all cases higher than the Ho value. Inbreeding coefficient per locus (F_is_) was positive for all markers and ranged from 0.071 to 1.000 (mean 0.323). Taken together, these findings indicate a low degree of homozygosity in the investigated samples.

**Table 2.** Genetic characteristics and variation of the 15 microsatellite loci detected in the population of dermotropic *L. infantum* strains (Emilia-Romagna region, northern Italy, n=44).

| Locus | n | Repeat array<br>per locus | Fragment size<br>array per locus (bp) | A | He | Ho | $F_{is}$ |
| --- | --- | --- | --- | --- | --- | --- | --- |
| Li41-56 | 44 | CA (7-11) | 86-94 | 4 | 0.731 | 0.568 | 0.224 |
| Li46-67 | 44 | CA (7-10) | 74-80 | 3 | 0.473 | 0.000 | 1.000 |
| Li21-34 | 44 | CA (8-15) | 81-95 | 4 | 0.403 | 0.205 | 0.495 |
| Li22-35 | 43 | CA (8-17) | 84-102 | 7 | 0.746 | 0.559 | 0.254 |
| Li23-41 | 44 | GT (12-19) | 77-91 | 7 | 0.744 | 0.523 | 0.300 |
| Lm2TG | 44 | TG (9-26) | 110-144 | 10 | 0.764 | 0.568 | 0.258 |
| Lm4TA | 44 | TA (9-20) | 73-95 | 11 | 0.764 | 0.545 | 0.288 |
| Li71-5/2 | 44 | CA (8-10) | 108-112 | 3 | 0.662 | 0.568 | 0.143 |
| LIST7039 | 44 | CA (11-18) | 199-213 | 8 | 0.821 | 0.318 | 0.615 |
| Li71-33 | 43 | TG (10-14) | 103-111 | 4 | 0.684 | 0.558 | 0.186 |
| Li71-7 | 44 | CA (7-13) | 88-100 | 6 | 0.792 | 0.614 | 0.227 |
| CS20 | 44 | TG (10-23) | 65-91 | 9 | 0.804 | 0.477 | 0.409 |
| Li45-24 | 42 | CA (7-17) | 89-109 | 8 | 0.694 | 0.476 | 0.316 |
| TubCA | 44 | CA (9-16) | 80-94 | 5 | 0.648 | 0.477 | 0.266 |
| LIST7031 | 43 | CA (8-12) | 165-173 | 5 | 0.601 | 0.558 | 0.071 |
| **Overall** | **43.7** |  |  | **6.3** | **0.689** | **0.468** | **0.323** |

Locus, analyzed microsatellite marker; n, sample size; Repeat array per locus, in brackets the overall identified dinucleotide repeat numbers; CA, dinucleotide repeat made of cytosine and adenine; TG, dinucleotide repeat made of thymine and guanine; Fragment size array per locus, the overall fragment sizes are indicated; bp, base pair; A, number of alleles; He, expected heterozygosity; Ho, observed heterozygosity; $F_{is}$ , inbreeding coefficient.

In total, 43 different MLG were assigned to 44 *L. infantum* samples (S1 Table). The only shared MLG was observed between strains ID6 and ID17 (Table 1 and S1 Table). The heterozygous allele combination predominated in the examined samples; five out of 44 were mono-allelic in all loci (ID12, ID22, ID33, ID40, ID44) and were deemed as homozygous.

### Inference of population structure

Given the observed genetic variability and the presence of distinct MLG, the integrated dataset was further investigated for evidence of population structure. The microsatellite profiles obtained for the 44 TL strains were compared with those of strains from different sources, including both previously characterized strains (13) and additional strains analyzed for the first time in this study, as detailed in the Materials and Methods section. The resulting integrated dataset revealed several shared MLG, both within and across sample groups (S1 Table), with clonal patterns prevailing among VL, sand fly and canine strains. Notably, the two TL strains ID22 and ID44 shared MLG21 and MLG43 with VL strains. To minimize bias from clonal redundancy, clone-correction was applied, retaining each unique MLG only once in subsequent analyses (N=106). STRUCTURE analysis indicated the presence of three distinct populations (Fig.1, S1 Table and S2 Table) as evidenced by ΔK values (S1 Figure).

**Figure 1.**
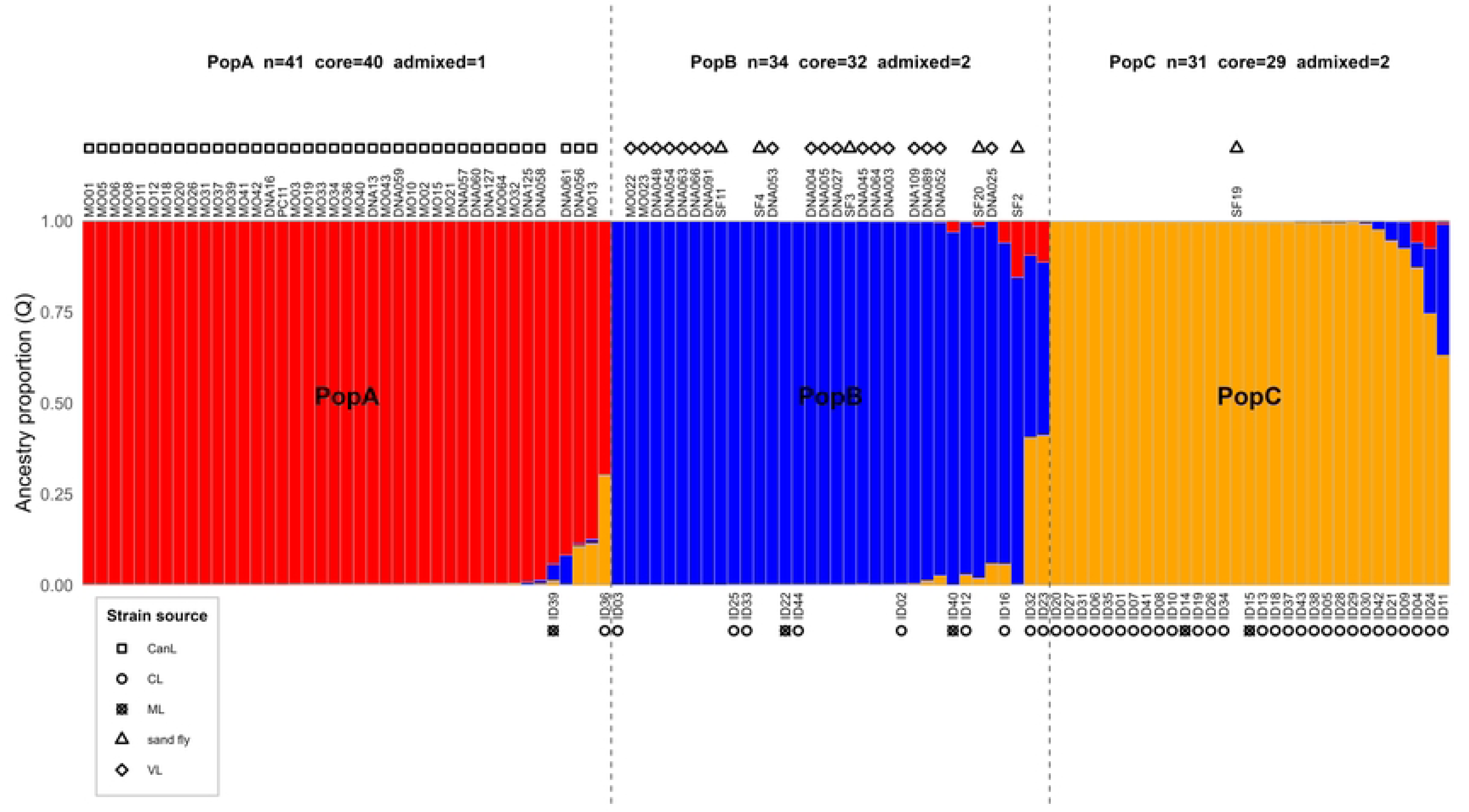
Estimated population structure for the *L. infantum* strains (dataset-2 clone-corrected; N=106) included in this study, as inferred by the STRUCTURE software on the basis of data for 15 microsatellite markers. Each strain is represented by a single vertical line divided into K colors, where K is the number of populations that were assumed. Each color represents one population, and the length of the colored segment shows the strain’s estimated proportion of membership (Q) in that population.

PopA included 40 canine strains and one TL strain (ID39 obtained from a patient residing in Apulia, southern Italy). PopB included 9 TL strains, all VL cases and five sand fly samples, while PopC included 28 TL strains and a strain from sand flies (S1 Table). Five TL strains, i.e., ID11, ID23, ID24, ID32, ID36 showed shared membership, with four displaying admixture between PopB and PopC, and one between PopA and PopC (Table 1).

Of the three strains collected outside E-R, one was assigned to PopA (ID39), one exhibited admixed membership between PopA and PopC, with a high membership coefficient for PopA (ID36), and one was admixed between PopB and PopC (ID24). Among the group with non-defined geographic origin, one strain was assigned to PopB (ID22), one to PopC (ID21), and two were classified as admixed PopB/PopC (ID23 and ID32).

The spatial distribution of the strains across the Bayesian-inferred genetic clusters is shown in Figure 2. Assignment to PopB or PopC of TL samples was independent of geographic origin within E-R. Furthermore, no temporal clustering was observed by year of collection (S1 Table), nor was there any association between the identified populations and the clinical form of TL (CL vs ML).

**Figure 2.**
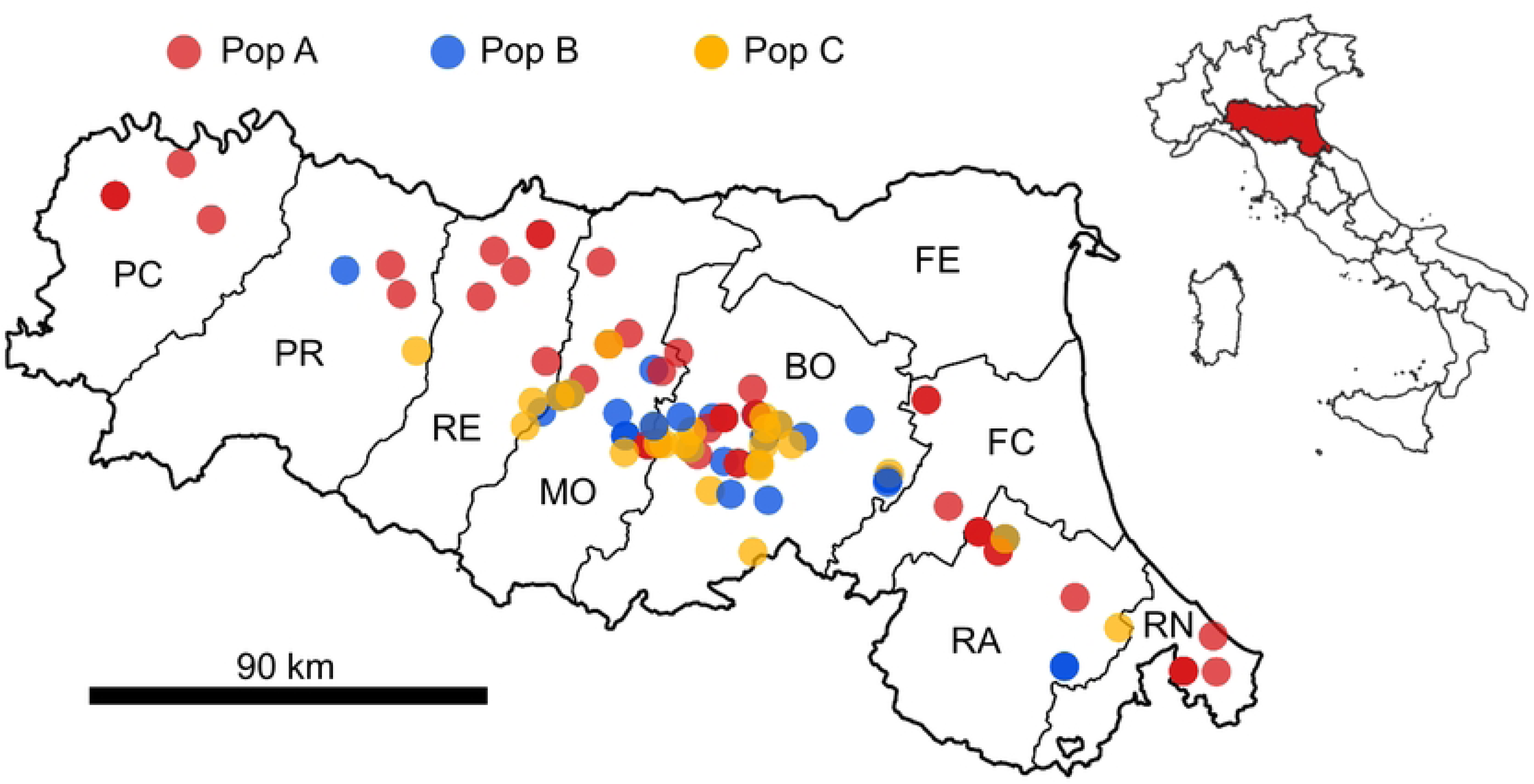
Geographical distribution of human (including cases of tegumentary and visceral leishmaniasis), canine and sand fly *Leishmania*-positive samples, 2013-2024, Emilia-Romagna region (northern Italy). Colors depict the populations inferred by STRUCTURE (K=3). Map generated with Quantum-GIS (https://www.qgis.org/it/site/).

Finally, following the STRUCTURE analysis performed on the clone-corrected dataset, the clones were reintroduced and assigned to the same population as their corresponding MLG. As a result, seven and six shared MLG were detected in PopA and PopB, respectively, and one in PopC (S1 Table).

To further explore potential substructure, each population was subsequently analyzed separately using the same Bayesian clustering framework. Within PopB, STRUCTURE consistently supported the presence of two clearly differentiated genetic groups, which did not show any clear association with geographic origin. A secondary peak at K=3 was observed, at which a more localized clustering emerged, involving a small subset of strains from the same area. For the PopA, the analysis identified a more complex pattern. While higher K values resulted in additional clusters, these were unstable across runs and involved limited numbers of strains and a simpler subdivision involving two major genetic groups represents the most parsimonious interpretation of the data. In contrast, for PopC no additional subdivision was detected (see S1 text for additional details).

### Multivariate analyses

Multivariate analyses were performed on the clone-corrected dataset (n=106) and were congruent with the population structure inferred by STRUCTURE. PCA revealed a clear separation of strains into three main genetic groups corresponding to PopA, PopB and PopC along the first two principal components, which accounted for a substantial proportion of the total genetic variance (Fig. 3A). PopC formed a distinct cluster in the multivariate space, whereas a small number of strains occupied intermediate positions between PopB and PopC. DAPC, performed using clusters inferred through a BIC-based approach without a priori population assignment, further supported the separation among the three populations (Fig. 3B). PopC formed a homogeneous cluster with no evidence of further subdivision, whereas PopA and PopB exhibited internal genetic structure. Most strains inferred as admixed by STRUCTURE were positioned closer to PopC in the discriminant space, with one exception (ID39) grouping within PopA. Additional details on multivariate analyses, including parameter optimization and exploratory clustering, are provided in S1 Text.

**Figure 3.**
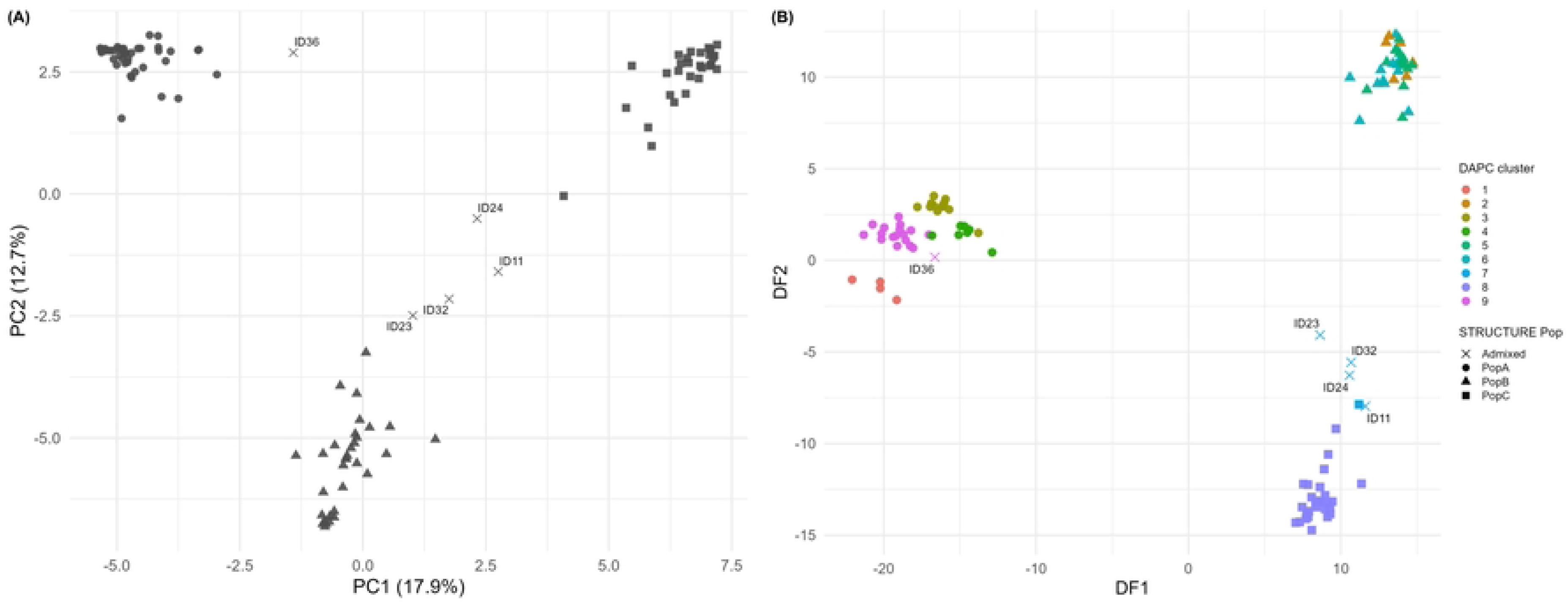
Multivariate analyses of *L. infantum* strains based on multilocus microsatellite typing (MLMT) data. (A) Principal component analysis (PCA) showing the distribution of individual genotypes along the first two principal components. (B) Unsupervised discriminant analysis of principal components (DAPC) based on 30 retained principal components and two discriminant functions. Colors represent genetic clusters inferred by DAPC, while point shapes indicate population assignment inferred by STRUCTURE.

### Genetic characterization of identified populations

For population genetic analyses, admixed strains were excluded (final N=101). Concordance across methods was further supported by population differentiation: pairwise fixation index (F_st_) values were above 0.25, indicating strong genetic differentiation between all main populations (Tab. 3).

**Table 3.**
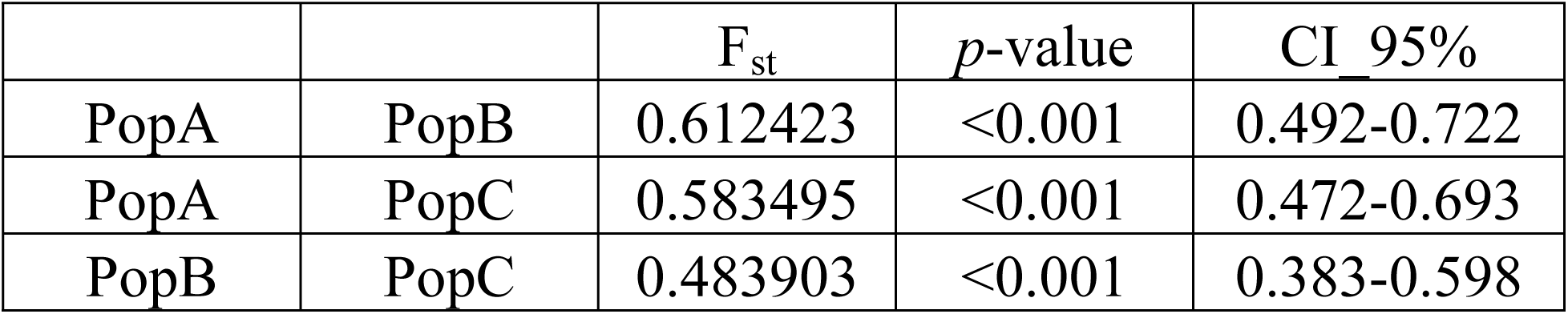
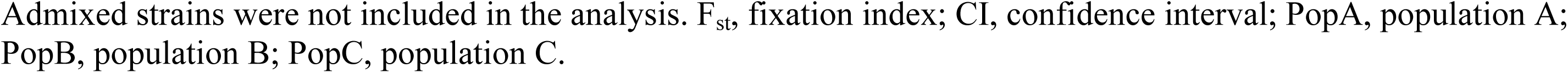
Differentiation measures (F_st_) and corresponding *p*-values for the populations and subpopulations of *L. infantum* strains (n=101) from the Emilia-Romagna region (northern Italy) as assumed by STRUCTURE.

| | | $F_{st}$ | $p$-value | CI 95% |
| --- | --- | --- | --- | --- |
| PopA | PopB | 0.612423 | <0.001 | 0.492-0.722 |
| PopA | PopC | 0.583495 | <0.001 | 0.472-0.693 |
| PopB | PopC | 0.483903 | <0.001 | 0.383-0.598 |

Admixed strains were not included in the analysis. $F_{st}$ , fixation index; CI, confidence interval; PopA, population A; PopB, population B; PopC, population C.

Descriptive analyses were conducted for each main population (Tab. 4). Tests for HWE revealed a significant departure from equilibrium across all populations (p<0.05), with many loci remaining in disequilibrium even after Bonferroni correction (S3 Table). However, the direction of deviation varied between populations. PopA and PopB consistently exhibited high and positive F_is_ values, reflecting a marked deficit of heterozygosity. This trend was mirrored by lower Ho values in both populations. On the other hand, PopC displayed a negative F_is_ (−0.197), indicating an excess of heterozygotes, and a higher genetic diversity, with MNA = 3.933 and He = 0.540.

**Table 4.**
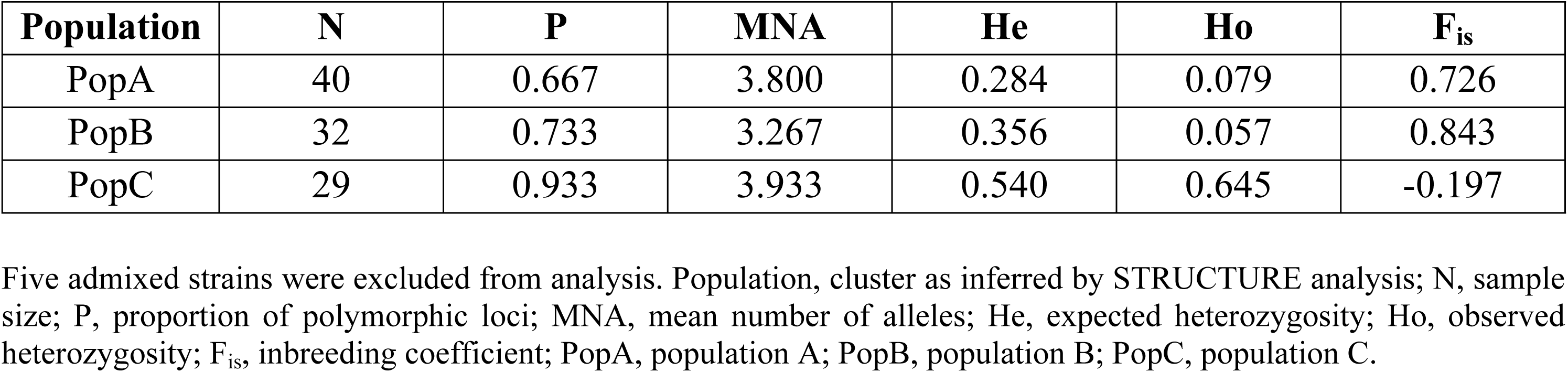
Genetic characterization of the *L. infantum* populations circulating in the Emilia-Romagna region, northern Italy (n=106).

| Population | N | P | MNA | He | Ho | $F_{is}$ |
| --- | --- | --- | --- | --- | --- | --- |
| PopA | 40 | 0.667 | 3.800 | 0.284 | 0.079 | 0.726 |
| PopB | 32 | 0.733 | 3.267 | 0.356 | 0.057 | 0.843 |
| PopC | 29 | 0.933 | 3.933 | 0.540 | 0.645 | -0.197 |

Five admixed strains were excluded from analysis. Population, cluster as inferred by STRUCTURE analysis; N, sample size; P, proportion of polymorphic loci; MNA, mean number of alleles; He, expected heterozygosity; Ho, observed heterozygosity; $F_{is}$ , inbreeding coefficient; PopA, population A; PopB, population B; PopC, population C.

### Microsatellite-based tree analysis

Microsatellite-based tree reconstruction (Fig. 4) yielded a topology consistent with the population structure inferred by STRUCTURE and DAPC. Three major clades corresponding to PopA, PopB and PopC were observed. The clade corresponding to PopA was composed mainly of canine strains and included the *L. infantum* reference strains representative of zymodemes MON-1 and MON-72. PopB formed a distinct clade comprising all VL-associated strains together with a subset of TL strains and sand fly samples. PopC formed a separate clade largely composed of TL-associated strains. The *L. donovani* reference strain formed a distinct, long branch positioned closer to PopB. Four of the five TL strains with admixed ancestry (ID11, ID23, ID24, ID32) were located outside the three main clades and clustered separately with the *L. infantum* MON-24 reference strains, while the remaining admixed strain (ID36) clustered together with ID39 within PopA.

**Figure 4.**
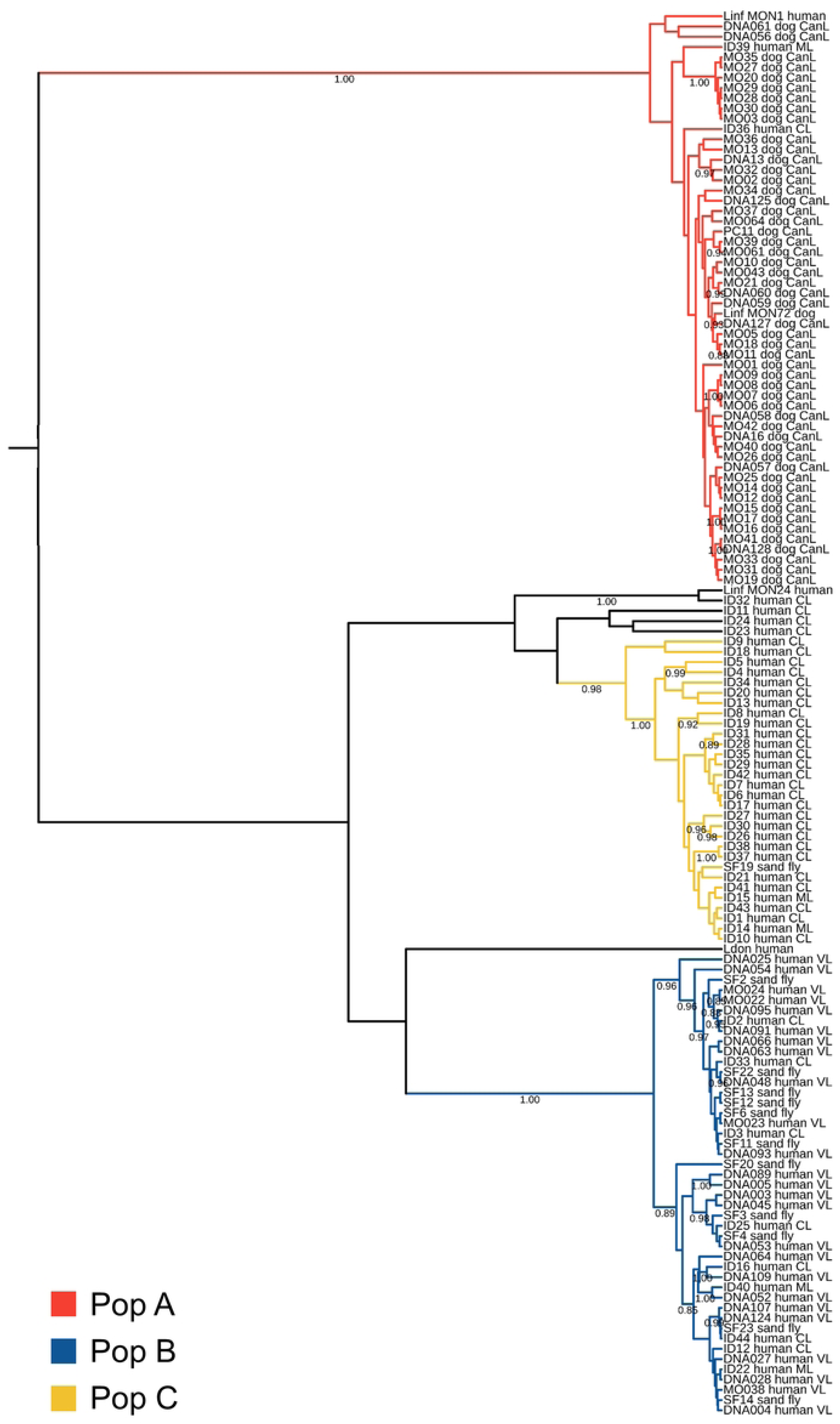
Microsatellite-based Bayesian tree inferred from multilocus microsatellite typing (MLMT) data using the Sainudiin model for the *Leishmania infantum* strains from this study (n=133) and n=4 WHO reference strains. Tree branches are colored according to population assignments inferred by STRUCTURE (K=3; PopA, red; PopB, blue; PopC, yellow). Posterior probability values > 0.85 are showed at the corresponding nodes. Representative strains for each multilocus genotype are listed in S1 Table. Strains labels specify the laboratory code, host of origin, and clinical form (CL, cutaneous leishmaniasis; ML, mucosal leishmaniasis; VL, visceral leishmaniasis; CanL, canine leishmaniasis). WHO reference strains are labelled by species abbreviation (Linf, *Leishmania infantum*; Ldon, *Leishmania donovani*) and host of origin.

## Discussion

Over the last decades, dermotropic strains of *L. infantum* have been increasingly reported through the Mediterranean basin, with a progressive geographical expansion and a rising number of cases (5). In E-R, numerous cases of CL were reported during the 1950s (35); after several decades of sporadically reported cases, a re-emergence of CL has been observed since 2010, concomitant with a rise of VL cases (8). This epidemiological scenario highlights the need to better characterize the population structure of *L. infantum* and its possible association with different clinical presentations.

MLMT revealed high genetic polymorphism among *L. infantum* strains associated with TL in northern Italy, as evidenced by the identification of 43 distinct MLG among the 44 analysed strains. When compared with sympatric strains from VL cases, dogs, and sand flies, the TL strains segregated into two main genetic populations, called PopB and PopC. PopB clustered together with strains associated with VL, whereas PopC was detected exclusively among TL cases. Notably, neither of these human-associated *L. infantum* populations overlapped with PopA, which comprises the most widespread *L. infantum* MON-1 group circulating in the Mediterranean basin. Only two TL strains clustered within PopA; both derived from infections acquired outside E-R (Liguria and Apulia, respectively), in line with our previous observations that human cases in E-R are caused by parasite populations not yet detected in autochthonous dogs (15).

Our findings reveal that PopB, previously associated with VL in E-R (13), also includes nine TL strains, suggesting that parasite genetic background alone may be insufficient to fully explain tissue tropism and that host-related factors play a major role in progression to systemic or localized disease (36). On the other hand, the identification of PopC represents a relevant novelty of this study. This population was strongly associated with TL cases, as most core PopC strains were derived from tegumentary infections. The presence of strains with predominant PopC ancestry (e.g., PopB/CC profiles in STRUCTURE), including strains from patients residing outside northern Italy (i.e., ID24), further supports a link between this genetic background and dermotropic manifestations. The distinct placement of these admixed strains in the microsatellite-based tree, outside the compact PopC clade, is consistent with their partial ancestry and highlights the dynamic genetic exchange among *L. infantum* populations in Italy. The diversity of TL-associated strains observed here is consistent with previous findings based on *Hsp*70 coding gene sequencing, supporting the complex population structure of *L. infantum* in this region (12,13).

High pairwise F_st_ estimates between the three populations indicated strong genetic differentiation, although strains with admixed ancestry were detected between PopB and PopC. The absence of any significant geographical or temporal clustering suggests that these populations co-circulate within the same endemic area, while remaining genetically distinct. Such segregation may reflect partially distinct transmission cycles or ecological niches, potentially involving different combinations of reservoirs and vectors. In this regard, reports about wild animals infected with *L. infantum* are increasing in the Mediterranean region (37) and in E-R (38). Moreover, PopB and PopC have been detected in sand flies in both the present and previous studies (13,15). While *P. perfiliewi* is the predominant sand fly species in E-R (39), the sympatric presence of *P. perniciosus*, even at a lower number, raises the hypothesis that the distinct vector-parasite interactions may contribute to maintaining population structure.

Marked differences in heterozygosity patterns were observed among populations. High F_is_ values observed in PopA and PopB align with previous reports in *L. infantum* and likely reflect combination of clonal propagation, inbreeding and microgeographic population structure, rather than a single underlying process (40). The presence of multiple shared MLG within these populations supports the existence of clonal expansion, which can contribute to deviations from HWE.

Similar patterns have been interpreted in *Leishmania* populations as the result of population subdivision and localized transmission foci, where clonality alternates with endogamic sexual reproduction (inbreeding) (41). Conversely, PopC exhibited a moderately negative mean F_is_ (-0.197), consistent with an excess of heterozygotes.

Such pattern is commonly observed in clonal organisms, where the accumulation of mutations leads to increased allelic divergence (42). However, the simultaneous presence of loci showing both positive and negative deviations from HWE is also compatible with partially clonal organisms, in which F_is_ values can vary within the same population depending on the relative contribution of genetic drift, mutation, and occasional sexual reproduction (43).

As an alternative explanation, the excess of heterozygosity and the lower number of shared MLG compared to PopA and PopB could reflect distinct demographic processes, with TL strains potentially experiencing weaker bottlenecks than VL strains, which on the other hand may face stronger immune pressure. Consistent with this hypothesis, negative F_is_ values have been reported in dermotropic *L. tropica* populations (44,45) and in *L. infantum* MON-24 strains isolated from CL cases (46). Overall, differences in F_is_ values among populations should be interpreted as signatures of distinct evolutionary and demographic trajectories rather than as direct indicators of current reproductive mode.

A major strength of this study is the One-Health design, integrating sympatric *L. infantum* strains from humans, dogs and sand flies collected in the same endemic area and over a comparable time frame. The concordant population inference across STRUCTURE, DAPC, Fst estimates and phylogenetic analysis, provided robust evidence for the existence of three highly differentiated populations and enabled the identification of a TL-associated lineage (PopC).

Some limitations should nevertheless be acknowledged. The number of TL cases was limited and geographically uneven, and sand fly sampling remained insufficient, restricting inferences on vector–parasite associations. Finally, microsatellites capture mainly neutral variation and do not allow direct inference on functional determinants of tissue tropism; therefore, the association of PopC with TL should be regarded as epidemiological and warrants validation through genome-wide and functional approaches.

### Conclusions

The analysis of *L. infantum* strains from TL cases revealed substantial genetic diversity among parasites circulating in northern Italy and led to the identification of a genetic population (PopC) detected exclusively among CL and ML cases in the study area. On the other hand, PopB encompassed strains from both TL and VL cases, supporting its role as a lineage with broad ecological adaptability. Finally, PopA, which is mainly composed of canine isolates, remained clearly separated from *L. infantum* populations that infect humans in northern Italy. The strong genetic differentiation observed among populations supports the existence of distinct parasite lineages that may follow heterogenous evolutionary and epidemiological trajectories. Overall, these findings highlight the need for integrated One-Health surveillance, combining highly discriminatory typing of sympatric strains from humans, vectors and animal hosts, to better clarify the population dynamics and epidemiology of leishmaniasis in Italy.

## Funding Acquisition

This study was supported by EU funding within the NextGeneration EUMUR PNRR Extended Partnership initiative on Emerging Infectious Diseases (Project no. PE00000007, INF-ACT) (SV), by the Ministry of Health, Italy (grant H77G23000320001 – IZS SI PRC 06/23) (FB) and by the Ministry of Health, Italy (grant E53C2200224000– IZS LER PRC 08/22/) (MC). The funders had no role in study design, data collection and analysis, decision to publish, or preparation of the manuscript.

## Acknowledgements

The authors gratefully acknowledge the Emilia-Romagna Region and the coordination team of the Regional Surveillance Plan for canine leishmaniasis, whose support made it possible to maintain the integrated surveillance framework on which this study is based. The authors also wish to thank Prof. Francesco Dondi and Dr. Francesco Lunetta (Department of Veterinary Medical Sciences, University of Bologna, Italy) for their collaboration and for providing two canine isolates included in this study, originally obtained during an independent project on drug resistance.

The Skin_Leish_RER network is composed by:

Maria Pia Foschini^1^, Iria Neri^2^, Cosimo Misciali^2^; Giulia Querzoli^3^, Barbara Corti^3^, Giacomo Nigrisoli^4^, Federica Lanna^4^, Bianca Granozzi^5^, Laura Gerna^6^, Rosa Scotti^7^, Anna Maria Cesinaro^7,8^, Erica Franceschini^9,10^, Simonetta Piana^11^, Vito Di Lernia^12^, Alberico Motolese^12^, Simona Schivazzappa^13^, Silvana Trincone^14^, Francesca Scarpellini^15^, Michela Tabanelli^16^, Paola Sgubbi^16^, Silvia Zago^17^, Luca Saragoni^4,17^, Romina Gianfreda^18^, Anna Rita Lombardi^19^, Antonella Frassetto^20^, Alessandro Borghi^21^.

^1^Department of Biomedical and Neuromotor Sciences, University of Bologna, Pathology Unit at Bellaria Hospital, Bologna, Italy.

^2^Unit of Dermatology, IRCCS Azienda Ospedaliero-Universitaria di Bologna, Bologna, Italy

^3^Pathology Unit, IRCCS Azienda Ospedaliero-Universitaria di Bologna, Bologna, Italy

^4^Department of Medical and Surgical Sciences, University of Bologna, Bologna, Italy

^5^Infectious Diseases Unit, IRCSS Azienda Ospedaliero Universitaria di Bologna, Bologna, Italy

^6^Infectious Diseases Unit, Guglielmo da Saliceto Hospital, Piacenza, Italy

^7^Pathology Unit, Azienda Ospedaliero-Universitaria di Modena, Modena, Italy

^8^Laboratorio Test s.r.l., Modena, Italy

^9^Infectious Diseases Unit, Azienda Ospedaliero-Universitaria di Modena, Modena, Italy

^10^Department of Surgery, Medicine, Dental Medicine and Morphological Sciences, University of Modena and Reggio Emilia, Modena, Italy

^11^Pathology Unit, Azienda USL-IRCCS Reggio Emilia, Reggio Emilia, Italy

^12^Dermatology Unit, Azienda USL-IRCCS Reggio Emilia, Reggio Emilia, Italy

^13^Unit of Infectious Diseases, Azienda Ospedaliero-Universitaria di Parma, Parma, Italy

^14^Dermatology and Venereology Operating Unit, Bufalini Hospital, Cesena, Italy

^15^Pathology Unit, Bufalini Hospital, Cesena, Italy

^16^Dermatology Unit, Santa Maria Delle Croci Hospital of Ravenna, AUSL Romagna, Ravenna, Italy

^17^Pathology Unit, Azienda USL della Romagna, Ravenna Hospital, Italy

^18^Infectious Disease Unit, Infermi Hospital, AUSL Romagna, Rimini, Italy

^19^Pathology Unit, Infermi Hospital, AUSL Romagna, Rimini, Italy

^20^Dermatology Unit, Infermi Hospital, AUSL Romagna, Rimini, Italy

^21^Section of Dermatology and Infectious Diseases, Department of Medical Sciences, University of Ferrara, Ferrara, Italy

## Author contributions

GR: Conceptualization, Investigation, Data Curation, Formal Analysis, Visualization, Writing-Original Draft Preparation

EC: Conceptualization, Investigation, Data Curation, Formal Analysis, Writing-Original Draft Preparation

MC: Resources, Investigation, Formal Analysis, Funding Acquisition, Writing-Review & Editing AR: Investigation, Writing-Review & Editing

FB (Federica Bergamini): Investigation, Writing-Review & Editing TG: Investigation, Data Curation, Writing-Review & Editing

MO: Resources, Data Curation, Writing-Review & Editing TL: Resources, Writing-Review & Editing

VG: Resources, Data Curation, Writing-Review & Editing GC: Resources, Writing-Review & Editing

FB (Federica Bruno): Funding Acquisition, Project Administration, Supervision, Writing-Review & Editing

GFS: Supervision, Validation, Writing-Review & Editing

SV: Conceptualization, Data Curation, Funding Acquisition, Supervision, Writing-Original Draft Preparation

Skin_Leish_RER_network: Resources, Writing-Review & Editing

## Supporting information

**S1 text.** Supplementary results.

**S1 Figure.** Results inferred by STRUCTURE for the clone-corrected dataset-2 (n=106). A) Plot of the ΔK values per population (K) according to the Evanno method (see Materials and Methods), B) Plot of mean log likelihood of K and its variance per each K value.

**S1 Table.** Designation and species characteristics of *Leishmania infantum* strains obtained from tegumentary leishmaniasis (TL) cases (sheet 1, n=44), canine leishmaniasis, visceral leishmaniasis and sand flies (sheet 2, n=89), WHO *Leishmania* reference strains (sheet 3, n=4) analyzed in this study or used for comparison. CL, cutaneous leishmaniasis; ML mucosal leishmaniasis; VL, visceral leishmaniasis; CanL, canine leishmaniasis; E-R, Emilia-Romagna region; PR, Parma; LE, Lecce; Bo, Bologna; MO, Modena; PC, Piacenza; RN, Rimini; RE, Reggio Emilia; FC, Forlì-Cesena; RA, Ravenna; MLG, multilocus genotype; MLMT, multilocus microsatellite typing; A.N., accession number.

**S2 Table.** Panel of strains investigated in this study. Mean membership coefficient (Q) values and population assignment inferred by STRUCTURE at K=3, according to the criteria described in the Materials and Methods section.

**S3 Table.** Results of Hardy-Weinberg equilibrium (HWE) tests ordered by population and locus. *p*, *p*-value; se, standard error of the *p*-value; p_adj_Bonf, Bonferroni corrected p-value; F_is__Wc, inbreeding coefficient according to Weir and Cockerham (1984).

## References

1. Antonia AL, Ko DC. Leishmaniasis: A spectrum of diseases shaped by evolutionary pressures across diverse life cycle. Evol Med Public Health. 2020;2020(1):139–40.

2. Silva-Moreira AL, Serravite AM, Rios-Barros LV, de Menezes JPB, Horta MF, Castro-Gomes T. New insights into the life cycle, host cell tropism, and infection amplification of Leishmania spp. Infect Immun. 2025 Jul 8;93(7):e0012325.

3. Ghosh S, Nath S, Roy K, Karmakar S, Pal C. Leishmaniasis: Tissue Tropism in Relation to the Species Diversity. In 2023. p. 133–53.

4. Burza S, Croft SL, Boelaert M. Leishmaniasis. Lancet Lond Engl. 2018 Sep 15;392(10151):951– 70.

5. Ait Maatallah I, Akarid K, Lemrani M. Tissue tropism: Is it an intrinsic characteristic of Leishmania species? Acta Trop. 2022 Aug;232:106512.

6. Manual on case management and surveillance of the leishmaniases in the WHO European region.

7. McCall LI, Zhang WW, Matlashewski G. Determinants for the development of visceral leishmaniasis disease. PLoS Pathog. 2013 Jan;9(1):e1003053.

8. Todeschini R, Musti MA, Pandolfi P, Troncatti M, Baldini M, Resi D, et al. Re-emergence of human leishmaniasis in northern Italy, 2004 to 2022: a retrospective analysis. Euro Surveill Bull Eur Sur Mal Transm Eur Commun Dis Bull. 2024 Jan;29(4):2300190.

9. Franceschini E, Puzzolante C, Menozzi M, Rossi L, Bedini A, Orlando G, et al. Clinical and Microbiological Characteristics of Visceral Leishmaniasis Outbreak in a Northern Italian Nonendemic Area: A Retrospective Observational Study. BioMed Res Int. 2016;2016:6481028.

10. Gaspari V, Gritti T, Ortalli M, Santi A, Galletti G, Rossi A, et al. Tegumentary Leishmaniasis in Northeastern Italy from 2017 to 2020: A Neglected Public Health Issue. Int J Environ Res Public Health. 2022 Nov 30;19(23):16047.

11. Franssen SU, Durrant C, Stark O, Moser B, Downing T, Imamura H, et al. Global genome diversity of the Leishmania donovani complex. eLife. 2020 Mar 25;9:e51243.

12. Gritti T, Carra E, Van der Auwera G, Solana JC, Gaspari V, Trincone S, et al. Molecular Typing of Leishmania spp. Causing Tegumentary Leishmaniasis in Northeastern Italy, 2014-2020. Pathog Basel Switz. 2023 Dec 24;13(1):19.

13. Bruno F, Castelli G, Li B, Reale S, Carra E, Vitale F, et al. Genomic and epidemiological evidence for the emergence of a L. infantum/L. donovani hybrid with unusual epidemiology in northern Italy. mBio. 2024 Jul 17;15(7):e0099524.

14. Gritti T, Chicharro C, Carrillo E, Solana JC, Moreno J, Carra E, et al. Combination of Cpb-Hsp70 typing methods reveals genetic divergence between Leishmania infantum strains causing human tegumentary leishmaniasis in northern Italy and central Spain: a retrospective study. Infect Dis Poverty. 2025 May 26;14(1):41.

15. Rugna G, Carra E, Bergamini F, Calzolari M, Salvatore D, Corpus F, et al. Multilocus microsatellite typing (MLMT) reveals host-related population structure in Leishmania infantum from northeastern Italy. PLoS Negl Trop Dis. 2018 Jul;12(7):e0006595.

16. Aluru S, Hide M, Michel G, Bañuls AL, Marty P, Pomares C. Multilocus microsatellite typing of Leishmania and clinical applications: a review. Parasite Paris Fr. 2015;22:16.

17. Schönian G, Kuhls K, Mauricio IL. Molecular approaches for a better understanding of the epidemiology and population genetics of Leishmania. Parasitology. 2011 Apr;138(4):405–25.

18. Maggio 2023 22:22 UM 19. Bollettino Ufficiale della Regione Emilia-Romagna — (BURERT). [cited 2026 Feb 12]. Bollettino Ufficiale della Regione Emilia-Romagna — (BURERT). Available from: https://bur.regione.emilia-romagna.it

19. Pritchard JK, Stephens M, Donnelly P. Inference of Population Structure Using Multilocus Genotype Data. Genetics. 2000 Jun 1;155(2):945–59.

20. Jombart T, Devillard S, Balloux F. Discriminant analysis of principal components: a new method for the analysis of genetically structured populations. BMC Genet. 2010;11(1):94.

21. Evanno G, Regnaut S, Goudet J. Detecting the number of clusters of individuals using the software STRUCTURE: a simulation study. Mol Ecol. 2005 Jul;14(8):2611–20.

22. Jakobsson M, Rosenberg NA. CLUMPP: a cluster matching and permutation program for dealing with label switching and multimodality in analysis of population structure. Bioinformatics. 2007 Jul 15;23(14):1801–6.

23. Cortes S, Maurício IL, Kuhls K, Nunes M, Lopes C, Marcos M, et al. Genetic diversity evaluation on Portuguese Leishmania infantum strains by multilocus microsatellite typing. Infect Genet Evol. 2014 Aug;26:20–31.

24. Jombart T, Ahmed I. *adegenet 1.3-1*: new tools for the analysis of genome-wide SNP data. Bioinformatics. 2011 Nov 1;27(21):3070–1.

25. Jombart T. *adegenet*: a R package for the multivariate analysis of genetic markers. Bioinformatics. 2008 Jun 1;24(11):1403–5.

26. RStudio Team. RStudio: Integrated Development Environment for R [Internet]. Boston, MA; 2025. Available from: https://www.rstudio.com/

27. Wickham H. ggplot2: elegant graphics for data analysis. Second edition. Cham: Springer international publishing; 2016. 1 p. (Use R!).

28. Lewis P, Zaykin D. Genetic Data Analysis: Computer program for the analysis of allelic data. Version 1.0 (d16c) [Internet]. 2001. Available from: program distributed by the authors

29. Curtin University. GENEPOP on the Web [Internet]. [cited 2025 Sep 25]. Available from: https://genepop.curtin.edu.au

30. Guo SW, Thompson EA. Performing the Exact Test of Hardy-Weinberg Proportion for Multiple Alleles. Biometrics. 1992 Jun;48(2):361.

31. Weir BS, Cockerham CC. Estimating F-Statistics for the Analysis of Population Structure. Evolution. 1984 Nov;38(6):1358.

32. Wright S. Evolution and the genetics of populations, volume 4: variability within and among natural populations. Vol. 4. Chicago, IL: University of Chicago press; 1978.

33. Bouckaert R, Heled J, Kühnert D, Vaughan T, Wu CH, Xie D, et al. BEAST 2: A Software Platform for Bayesian Evolutionary Analysis. Prlic A, editor. PLoS Comput Biol. 2014 Apr 10;10(4):e1003537.

34. Sainudiin R, Durrett RT, Aquadro CF, Nielsen R. Microsatellite mutation models: insights from a comparison of humans and chimpanzees. Genetics. 2004 Sep;168(1):383–95.

35. Pampiglione, Silvio. La leishmaniosi viscerale in Emilia-Romagna. Ann Sanità Pubblica. 1974;34:1021–8.

36. Pratlong F, Rioux JA, Marty P, Faraut-Gambarelli F, Dereure J, Lanotte G, et al. Isoenzymatic Analysis of 712 Strains of *Leishmania infantum* in the South of France and Relationship of Enzymatic Polymorphism to Clinical and Epidemiological Features. J Clin Microbiol. 2004 Sep;42(9):4077–82.

37. Azami-Conesa I, Gómez-Muñoz MT, Martínez-Díaz RA. A Systematic Review (1990-2021) of Wild Animals Infected with Zoonotic Leishmania. Microorganisms. 2021 May 20;9(5):1101.

38. Taddei R, Bregoli A, Galletti G, Carra E, Fiorentini L, Fontana MC, et al. Wildlife Hosts of Leishmania infantum in a Re-Emerging Focus of Human Leishmaniasis, in Emilia-Romagna, Northeast Italy. Pathog Basel Switz. 2022 Nov 7;11(11):1308.

39. Calzolari M, Carra E, Rugna G, Bonilauri P, Bergamini F, Bellini R, et al. Isolation and Molecular Typing of Leishmania infantum from Phlebotomus perfiliewi in a Re-Emerging Focus of Leishmaniasis, Northeastern Italy. Microorganisms. 2019 Dec 3;7(12):644.

40. Kuhls K, Alam MZ, Cupolillo E, Ferreira GEM, Mauricio IL, Oddone R, et al. Comparative Microsatellite Typing of New World Leishmania infantum Reveals Low Heterogeneity among Populations and Its Recent Old World Origin. PLoS Negl Trop Dis. 2011 Jun 7;5(6):e1155.

41. Rougeron V, De Meeûs T, Hide M, Waleckx E, Bermudez H, Arevalo J, et al. Extreme inbreeding in Leishmania braziliensis. Proc Natl Acad Sci U S A. 2009 Jun 23;106(25):10224–9.

42. Prugnolle F, Meeûs TD. The impact of clonality on parasite population genetic structure. Parasite. 2008 Sep 1;15(3):455–7.

43. Reichel K, Masson JP, Malrieu F, Arnaud-Haond S, Stoeckel S. Rare sex or out of reach equilibrium? The dynamics of FIS in partially clonal organisms. BMC Genet. 2016 Jun 10;17:76.

44. Schwenkenbecher JM, Wirth T, Schnur LF, Jaffe CL, Schallig H, Al-Jawabreh A, et al. Microsatellite analysis reveals genetic structure of *Leishmania tropica*. Int J Parasitol. 2006 Feb 1;36(2):237–46.

45. Hakkour M, Badaoui B, El Hamiani Khatat S, Sahibi H, Fellah H, Sadak A, et al. Genetic diversity in Leishmania infantum and Leishmania tropica isolates from human and canine hosts in northern Morocco. Gene. 2024 Aug 30;921:148484.

46. Chargui N, Amro A, Haouas N, Schönian G, Babba H, Schmidt S, et al. Population structure of Tunisian *Leishmania infantum* and evidence for the existence of hybrids and gene flow between genetically different populations. Int J Parasitol. 2009 Jun 1;39(7):801–11.

